# A fast, reproducible, high-throughput variant calling workflow for evolutionary, ecological, and conservation genomics

**DOI:** 10.1101/2023.06.22.546168

**Authors:** Cade D. Mirchandani, Allison J. Shultz, Gregg W.C. Thomas, Sara J. Smith, Mara Baylis, Brian Arnold, Russ Corbett-Detig, Erik Enbody, Timothy B. Sackton

**Affiliations:** Department of Biomolecular Engineering, University of California, Santa Cruz; Santa Cruz, CA 95064, USA; Genomics Institute, University of California, Santa Cruz; Santa Cruz, CA 95064, USA; Ornithology Department, Natural History Museum of Los Angeles County, Los Angeles, CA 90007, USA; Informatics Group, Harvard University, Cambridge, MA, USA; Biology, Mount Royal University, Calgary, AB, T3E 6K6, CAN; Department of Computer Science, Princeton University, Princeton, NJ, USA; Center for Statistics and Machine Learning, Princeton University, Princeton, NJ, USA

**Author notes:** these authors contributed equally (Russ Corbett-Detig, Erik Enbody, Timothy B. Sackton).

## Abstract

The increasing availability of genomic resequencing datasets and high quality reference genomes across the tree of life present exciting opportunities for comparative population genomic studies. However, substantial challenges prevent the simple reuse of data across different studies and species, arising from variability in variant calling pipelines, data quality, and the need for computationally intensive reanalysis. Here, we present snpArcher, a flexible and highly efficient workflow designed for the analysis of genomic resequencing data in non-model organisms. snpArcher provides a standardized variant calling pipeline and includes modules for variant quality control, data visualization, variant filtering, and other downstream analysis.Implemented in Snakemake, snpArcher is user-friendly, reproducible, and designed to be compatible with HPC clusters and cloud environments. To demonstrate the flexibility of this pipeline, we applied snpArcher to 26 public resequencing datasets from non-mammalian vertebrates. These variant datasets are hosted publicly to enable future comparative population genomic analyses. With its extensibility and the availability of public datasets, snpArcher will contribute to a broader understanding of genetic variation across species by facilitating rapid use and reuse of large genomic datasets.

## Introduction

In the past decade, rapidly declining sequencing costs have led to a dramatic expansion in the availability of genomic resequencing datasets in diverse organisms, fueling a wide range of novel insights, including the prevalence of adaptive introgression between species (Huerta-Sánchez et al. 2014; Lamichhaney et al. 2015; Jones et al. 2018), the molecular basis of repeated local adaptation (Jones et al. 2012; Hill et al. 2019; Wooldridge et al. 2022), and the complex demographic histories of humans (Nielsen et al. 2017; Fan et al. 2023) and animals of conservation relevance (Robinson et al. 2018). In parallel, rapidly expanding efforts to generate high quality reference genomes across the Tree of Life (Rhie et al. 2021); (Lewin et al. 2022) are poised to empower population genetic inference across a wide diversity of organisms. The massive accumulation of existing genomic datasets facilitated by these advances can enable broad comparisons between diverse populations and uncover generalized principles that may explain processes that generate diversity across life. These questions include the determinants of molecular variation among species (Romiguier et al. 2014; Corbett-Detig et al. 2015; Buffalo 2021) and indirect estimates of the rates of loss of genetic variation among populations (Exposito-Alonso et al. 2022).

However, despite the rapid increase in accessibility of public sequencing data from diverse organisms, comparative population genetics and reuse of public data remains challenging for several reasons. In the absence of standardized variant calling pipelines for non-human species (Regier et al. 2018), computational batch effects introduced by differences in reference choice, alignment, and variant calling algorithms complicate efforts to jointly analyze existing variant calls across populations and species (Lek et al. 2016; Jia et al. 2020; Breton et al. 2021).

Considerations must also be given to data quality prior to data processing, particularly in cases of low coverage (Lou et al. 2021), and workflows must be flexible to accommodate these considerations. Because these computational and algorithmic choices can impact downstream analysis (Kulkarni and Frommolt 2017), comparative projects often must reanalyze raw data to produce comparable datasets, which can be computationally expensive.

Extensible, reproducible bioinformatic pipelines can help address these challenges, to facilitate both primary analysis of complex tasks such as variant calling and also allow for consistent reanalysis (Wratten et al. 2021). While reproducible workflows have had a major impact in human population genetics (Chen et al. 2022), the need for significant expertise to adapt pipelines optimized for human genetics to diverse species is a major technical hurdle for many researchers. Additionally, resequencing datasets are increasingly rapidly in scale (Ellegren 2014), driving a need for workflows optimized for computational efficiency and flexibility to use across a variety of compute resources, including cloud resources that eliminate the need for costly on-site infrastructure (Mangul et al. 2019).

Due to the popularity and need for efficient and reproducible workflows, several solutions have already been proposed for variant calling pipelines (Czech and Exposito-Alonso 2022; Cullen and Friedenberg 2023). Here, we present snpArcher, a reproducible workflow for dataset acquisition, variant calling, quality control, and downstream analysis that is optimized for nonmodel organisms and comparisons across datasets, available at https://github.com/harvardinformatics/snpArcher. snpArcher implements a combination of several notable features not included in other existing solutions that address the challenges presented by the expanding scale of comparative population genomics studies. First, our workflow is optimized for non-model species, which often lack gene annotations, known variant sites, and other genomic information typically required for human-optimized pipelines. Second, we take advantage of the huge compute power available through cloud resources and large high-performance computer (HPC) clusters by highly parallelizing the workflow’s variant calling step and thus greatly reducing analysis time. Third, we present a framework for extensibility and community development by defining downstream module contribution guidelines, including exemplary modules offering variant quality control, visualization, and other analyses.

To enable rapid analysis of a growing set of variant calls created in a functionally equivalent way, we apply this workflow to reanalyze public sequencing data and produce variant calls for 26 focal species of non-mammalian vertebrates, hosted for public use via Globus (accessible at https://app.globus.org/file-manager?origin_id=a6580c44-09fd-11ee-be16-195c41bc0be4&path=%2F or by a search for the “Comparative Population Genomics Data” public collection on Globus). Furthermore, we provide examples of analysis and visualization modules, and we use these to exemplify and enumerate a suite of criteria for future module contributions to this project. This new and immediately available toolset will enable highly reproducible comparative population genomic analyses for a broad range of taxa.

## Results/Discussion

### Overview of snpArcher

We developed snpArcher, a comprehensive workflow for the analysis of polymorphism data sampled from non-model organism populations (Figure 1). This workflow accepts raw sequence data and a reference genome as input, and ultimately produces a filtered, high quality VCF file for downstream analysis. This workflow is implemented in Snakemake (Mölder et al. 2021) and can be deployed on a HPC cluster or cloud environment to enhance scalability and accessibility. snpArcher uses a straightforward configuration file to flexibly accommodate many possible ways of running this pipeline. For example, it is possible to specify and download read data and reference genomes from NCBI or to specify local files for one or both of these inputs. snpArcher is designed to produce bioinformatics-standard data objects that can be used in a range of downstream applications. Furthermore, a defined module contribution model ensures that the workflow can be incorporated for use in a range of analyses and will grow through user contributions. snpArcher therefore provides a backbone for future development as well as a substrate for immediately comparable datasets.

**Figure 1.**
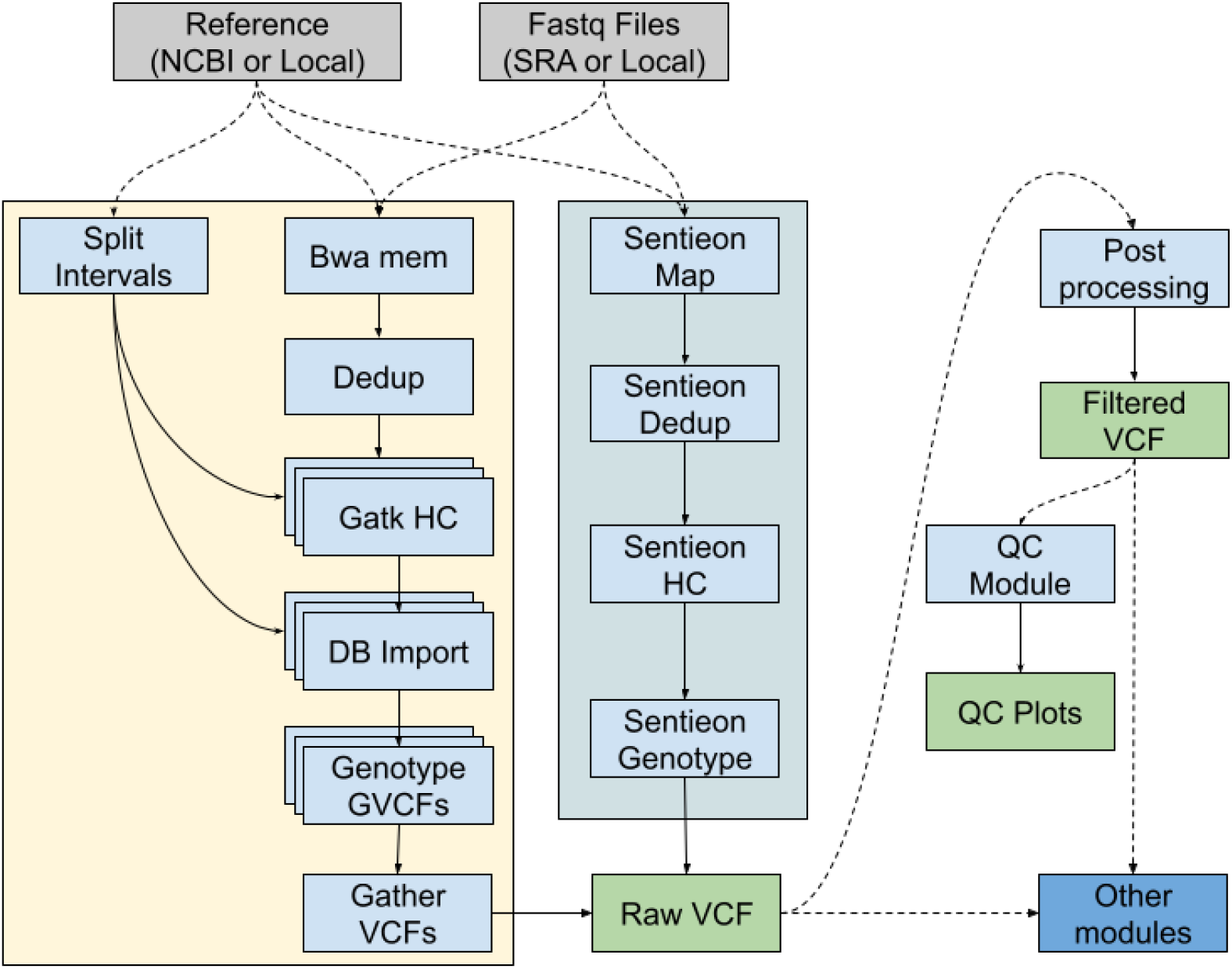
snpArcher overview. snpArcher is an automated pipeline implemented in Snakemake (Mölder et al. 2021). It takes short read whole genome sequencing data (fastq) and a reference genome as input and produces a multisample variant callset (VCF). With the modules presented here, snpArcher performs basic Quality Control statistics and visualizations.

### Datasets Processed and Public Database for Comparative Population Genomics

To thoroughly evaluate snpArcher and to provide a database of comparative population genomic datasets, we ran the workflow on 26 public resequencing datasets (Table S1). This dataset includes 13 birds, 12 fish, and 1 reptile. Datasets vary by number of individuals from 6 to 306, all with a mean depth of coverage of at least 5. Of these datasets, 13 are multispecies samples mapped to a common reference genome, 7 are primarily a single species but with one or two outgroup samples, and 6 are purely a single species.

A crucial advantage of snpArcher is that datasets will be maximally comparable across species because the bioinformatic processing is standardized. To ensure that future projects that use this pipeline have an immediately available set of comparable data, the VCFs and gVCFs for these datasets are hosted publicly on Globus (https://www.globus.org/)(Foster 2011; Allen et al. 2012) available at no cost to the user via the Comparative Population Genomics Data public collection. We expect that these datasets, in combination with the snpArcher pipeline, will spur future comparative population genomic analyses.

### Benchmarks

To evaluate the performance of snpArcher, we selected 10 individuals from a high quality resequencing dataset of Zebra finch (Singhal et al. 2015) and reanalyzed them using a range of approaches. First, we investigated the impacts of low sequencing depth by subsampling the initially high-depth dataset (16.7x-50.2x coverage) to uniform reduced coverage datasets (4x, 10x, and 20x). We ran each dataset using the “low coverage” and “high coverage” configurations of the pipeline; the “low coverage” configuration alters certain GATK parameters to improve single nucleotide polymorphism (SNP) calling in low coverage datasets. After filtering for SNPs that passed all filters, we genotyped about 40, 55, and 50 million SNPs in the 4x, 10x, and 20x datasets, respectively, with about 1 million more SNPs recovered from the low coverage pipeline at 4x coverage compared to the high coverage version. There were negligible differences for the two pipeline versions at 10x and 20x (Figure 2A). CPU time to run the low coverage version of the pipeline was substantially higher compared to the high depth version, and increased with sequencing depth (Figure 2B). The percentage of heterozygous sites per individual was substantially reduced at 4x coverages, the lowest with the 4x high depth pipeline, and slightly reduced at 10x coverage (Figure 2C). Individual fixation indices measuring expected heterozygosity (F statistics) were correspondingly higher at lower sequencing depths and with the high depth pipelines (Figure 2D), indicating less heterozygous dropout. While heterozygous dropout is a substantial problem at low-coverage (Nevado et al. 2014; Benjelloun et al. 2019), parameter tuning can partially mitigate its impact on genotype calls.

**Figure 2.**
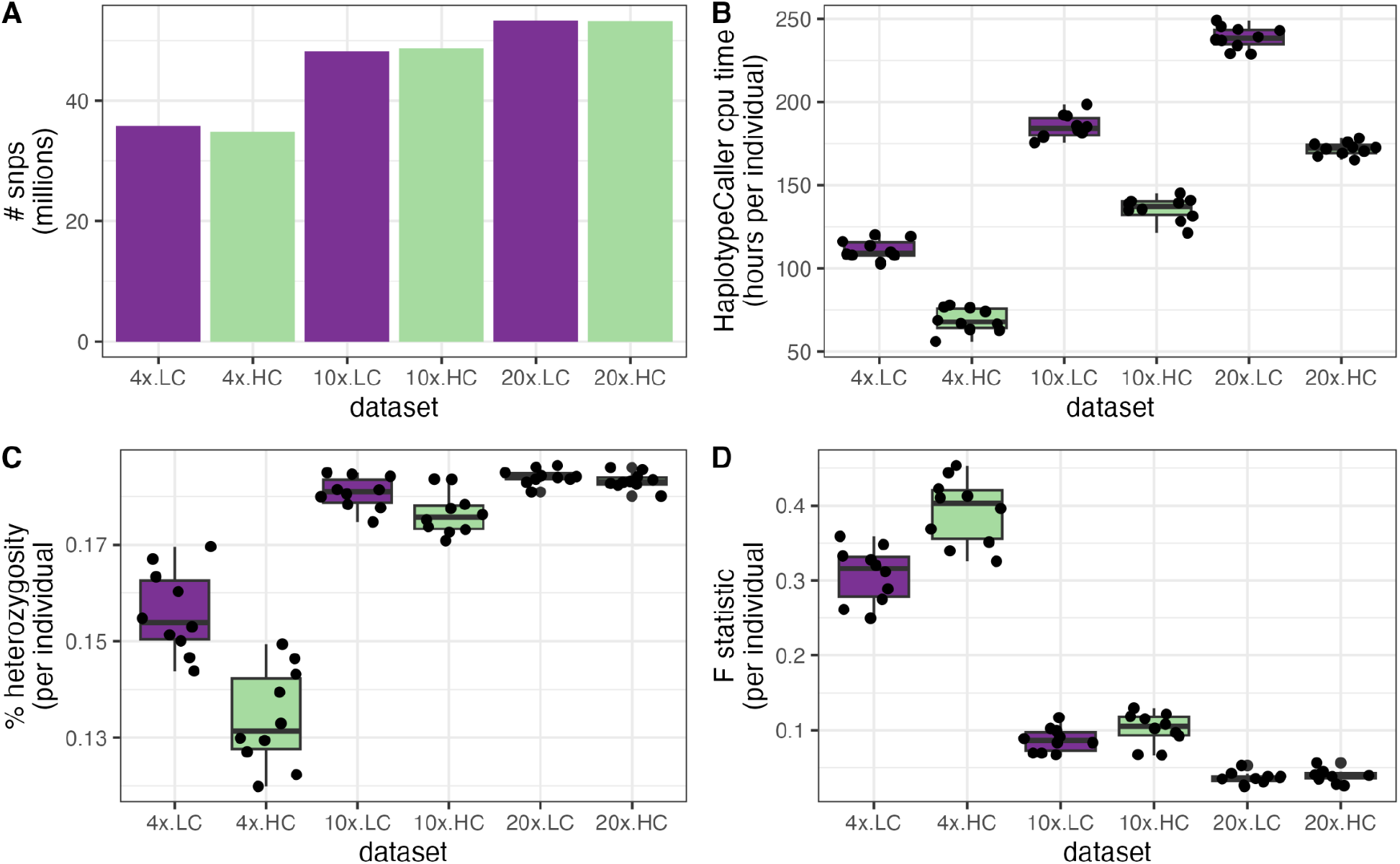
Benchmarks for the 10 individual Zebra Finch coverage and pipeline testing. For each coverage dataset (4x, 10x, 20x), we ran the low coverage (LC; purple) and high coverage (HC; green) version of the pipeline, and calculated A) the overall number of SNPs following standard SNP filtering, B) the hours of CPU time to run HaplotypeCaller for each individual, C) the percentage of heterozygous sites for each individual, and D) the F-statistic calculated for each individual.

Second, we assessed the effectiveness of our parallelization method for variant calling using snpArcher on the 10x Zebra finch dataset. A performance comparison was conducted between our scatter-by-Ns approach and the traditional scatter-by-chromosome approach. Given that GATK HaplotypeCaller has limitations in efficiently utilizing multiple CPU cores, it is recommended to parallelize this step using a scatter-gather technique, distributing the computation across the reference genome’s chromosomes (Heldenbrand et al. 2019). However, as runtime scales with genomic interval size (Fig 3A), using this approach will still result in potentially long execution times. To address this, we employ a strategy of partitioning chromosomes at Ns (assembly gaps), creating additional genomic intervals that enable further parallelization of the HaplotypeCaller step. This leads to increased compute utilization and reduced runtime per sample (Fig 3B). Although the effectiveness of this approach is dependent on available compute resources, the wide availability of high-performance computing (HPC) clusters and affordable cloud compute resources renders this constraint generally acceptable.

**Figure 3.**
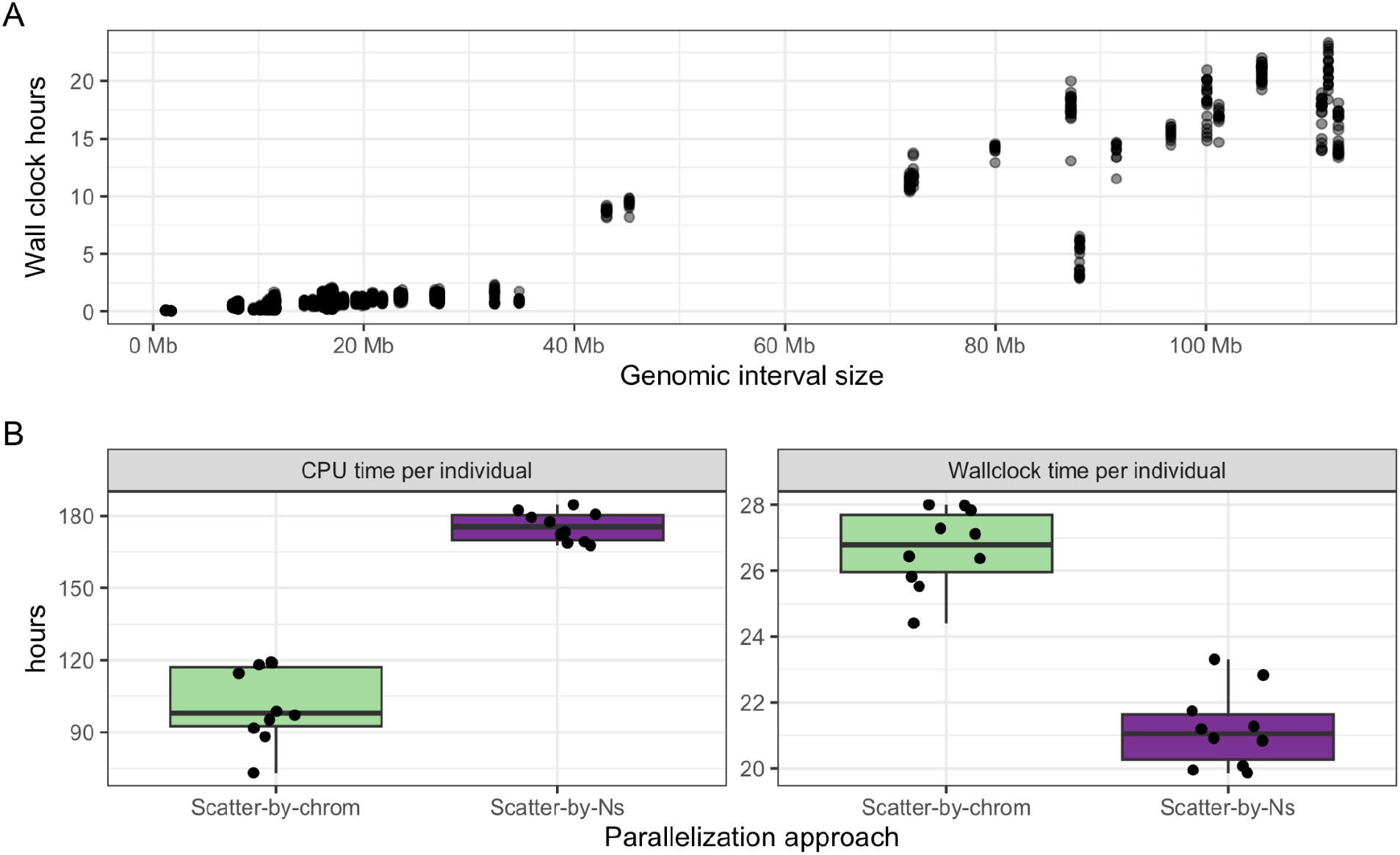
Wall clock and CPU time metrics from the HaplotypeCaller step of snpArcher. A) Wall clock time required to run HaplotypeCaller on genomic intervals. B) CPU time and wall clock time elapsed per individual to complete the HaplotypeCaller step using the scatter-by-chrom approach (green), or scatter-by-Ns approach (purple).

### Extensibility of snpArcher

A key goal in the design of the snpArcher pipeline is to allow seamless extensibility with downstream processing. We implement this using Snakemake modules, which allow additional rules to easily extend the main pipeline. To be added to snpArcher, a Snakemake workflow only needs a way to indicate that it should be run, such as a flag in the config file or a column in the sample sheet, and for output files from snpArcher to be linked to input files of the workflow. As long as these constraints are met, any user-defined Snakemake workflow can be imported as a module. We present several modular extensions of snpArcher here, but we hope also that user-developed modules will grow the set of tools linked to snpArcher in order to facilitate diverse analysis. We define a set of criteria for contributors to follow in the workflow documentation when adding additional modules.

### Quality Control and Data Visualization

An important component of any pipeline is quality control and data visualization outputs. We have implemented a module in snpArcher, run by default, that produces an interactive quality-control dashboard, which can be used to evaluate individual-level sequencing quality (Fig 4). This dashboard generates ten figures that allow visualization of basic summary statistics relating to population structure, batch effects, sequencing depth, genetic relatedness, geography, and admixture. For speed, most of these summaries are based on a random sample of 100,000 SNPs from across the genome. Four panels at the top of the dashboard provide high level summaries of the full variant dataset (i.e. without random downsampling to 100,000 SNPs).

**Figure 4.**
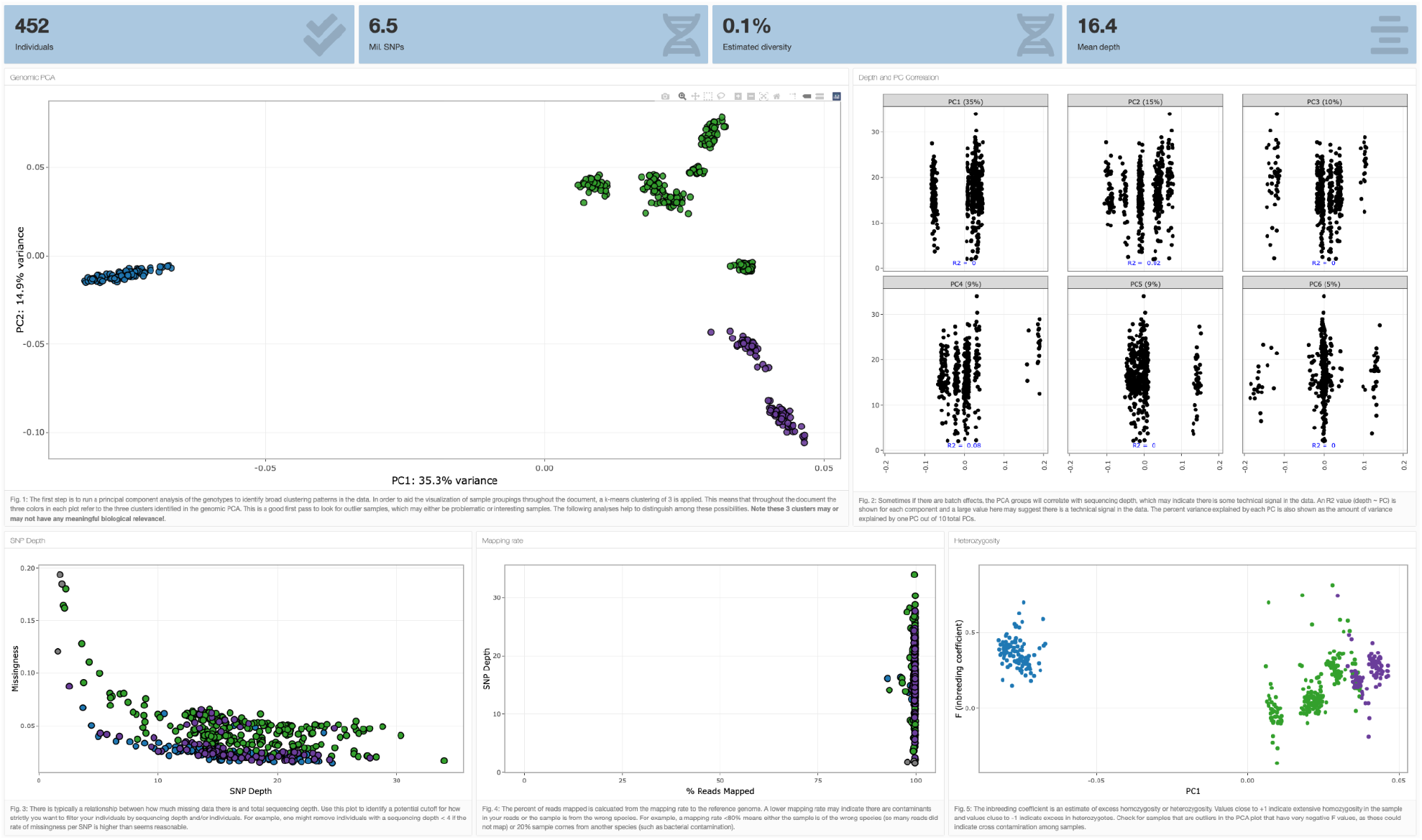
Preview of QC Dashboard for evaluating individual sequencing quality metrics. Shown here, genomic PCA, correlations between PCs and sequencing depth, relationship between missingness and SNP depth, percent mapped reads and SNP depth, and *FI* (inbreeding coefficient) and PC1. A complete interactive example can be found at [https://erikenbody.github.io/snpArcher/GCA_013435755.1_final_qc.html]

The use case for these simple visualizations is to quickly evaluate potential biases relating to individual-level sequencing variation. For example, in the principle component analysis (PCA) shown in the upper left panel of Fig. 4, it is possible to identify outliers that may represent cryptic genetic variation, batch effects, or otherwise problematic (or interesting) samples. By default, we identify three clusters based on PC1 and PC2 with k-means clustering (modifiable in the config file), and the remainder of the plots are colored according to these three clusters. Several metrics allow for the user to identify potential sequencing artifacts, for example by looking for associations between sequencing depth and PCA cluster (Fig. 4, upper right panel) or reference bias (Fig. 4, lower middle panel). An interactive heatmap of relatedness facilitates a rapid identification of close relatives in the dataset that may have otherwise been overlooked. Finally, two maps project spatial data as an interactive plot and provide a first pass visualization of the PCA clusters in space.

### Postprocessing

By default, snpArcher produces a raw variant call format (VCF) file with only basic filtering annotated. However, after viewing the individual-level quality-control visualizations as part of the QC module, users may wish to remove certain individuals from the analysis and apply additional filters on called variants. Additional postprocessing steps are implemented in a module, which runs if the user adds a column to the sample sheet header “SampleType.” The postprocessing module will exclude from the filtered VCF any sample with “exclude” as the SampleType, retaining all other individuals. Following this sample filtering, this module implements additional user-configurable filters. By default, the postprocessing workflow removes sites that fall into regions of low mappability, regions with excess coverage, and regions with insufficient coverage (defined by the configuration file), and then removes sites with a minor allele frequency < 0.1 or missingness > 75%. These thresholds can be configured by the user. Finally, two clean variant files are produced for SNPs and indels separately.

### MacDonald-Kreitman Tests

To demonstrate the potential to extend snpArcher to incorporate downstream analysis, we developed a module to evaluate positive selection among a sample of individuals from a population (the ingroup) as well as one or more diverged samples (the outgroup) by computing MacDonald-Kreitman (MK) tests for each gene (McDonald and Kreitman 1991). This module is triggered when samples are annotated as “ingroup” and “outgroup” using the SampleType column in the sample sheet. Samples that do not have either designation will be excluded from the MK tests.

To facilitate the development of this module, we wrote a stand-alone Python program, degenotate (https://github.com/harvardinformatics/degenotate), that can retrieve coding sequences from an annotated genome, compute degeneracy across the genome, and calculate MK tables; degenotate can be installed via conda and run independently, but is also incorporated into snpArcher’s MK module. Briefly, degenotate assesses whether SNPs in the postprocessed VCF encode for polymorphic sites within the ingroup or fixed differences between the ingroup and the outgroup. It further classifies whether each SNP, whether polymorphic or fixed, is synonymous or non-synonymous. Note that certain assumptions, detailed in the Methods, must be made about how to handle certain rare edge cases when doing this.

Based on these outputs, the MK module (or standalone degenotate) creates tables that are organized by gene and can be analyzed using the standard MacDonald-Kreitman test statistic, using various extensions (Rand and Kann 1996) (Stoletzki and Eyre-Walker 2011), or in aggregate to investigate genome wide signatures of natural selection (Messer and Petrov 2013). This module will enable rapid application of population-genomic tests of selection (Figure 5) and in combination with the database of processed population datasets, provides a framework for comparing rates of adaptation to a range of species.

**Figure 5.**
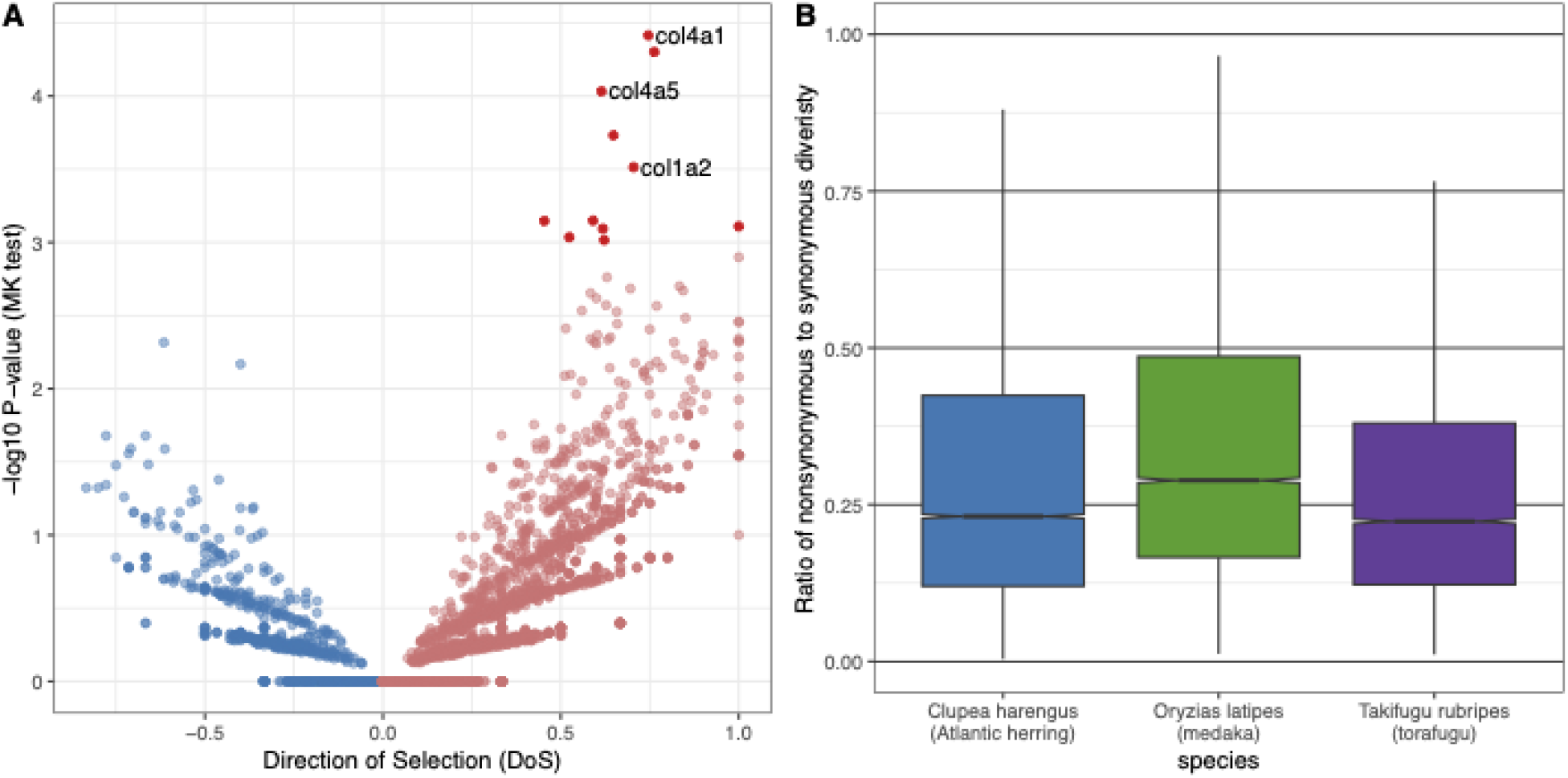
Analysis of three fish datasets using snpArcher+degenotate, demonstrating the possible applications of this module. **(A)** MK results for Takifugu rubripres, plotting the -log10 P-value of a Fisher’s exact test of the MK table on the Y axis, and the direction of selection on the X axis (Stoletzki and Eyre-Walker 2011; positive for an excess of nonsynonymous divergence, negative for an excess of nonsynomyous polymorphisms). Genes with nominal P-values < 0.001 are shaded darked, and three collagen genes with potential roles in tooth and spine development are highlighted. **(B)** Ratio of nonsynonymous to synonymous diversity (calculated based on number of segregating sites in each category) for three fish species. Boxplot show the median and interquartile range for protein coding genes in the genomes of each species.

### UCSC Genome Browser Track Data Hub Generation

To facilitate downstream data exploration and as an example of the module development components of this work, we developed a module to generate UCSC Genome Browser track files to explore population variation data (see Methods). Briefly, this module computes and generates genome browser tracks for traditional population genomic summary statistics such as windowed estimates of Tajima’s D, SNP density, Pi, Minor Allele Frequency, SNP depth. The Genome Browser tracks allow for rapid analysis of common population genomic statistics along with other available genomic feature tracks in an easy to access and shareable format (Fig 6).

**Figure 6.**
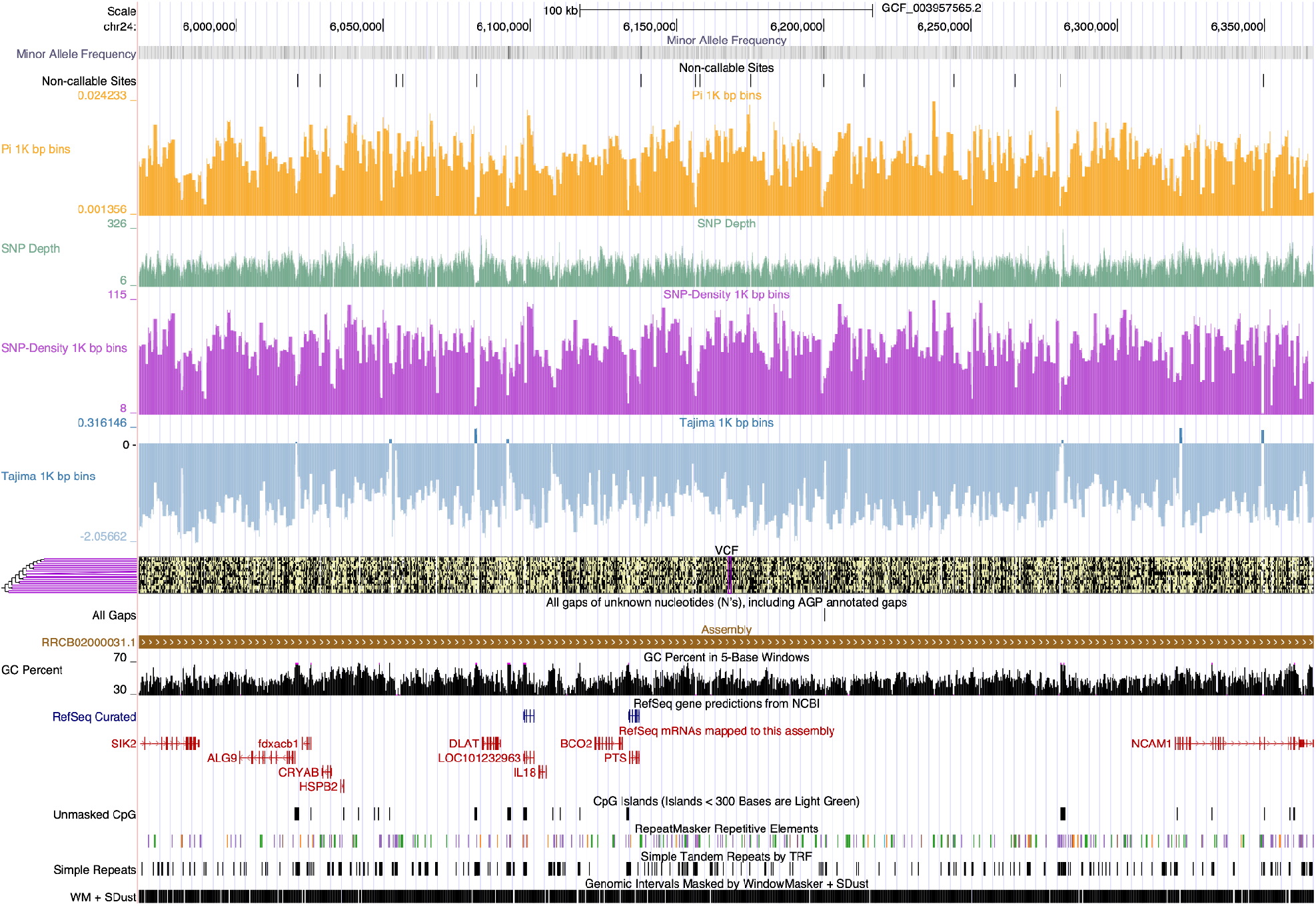
Example Genome Browser track hub created by the trackhub module. Tracks include: Minor allele frequency, non-callable sites, nucleotide diversity (Pi) in 1kb bins, SNP depth, SNP density in 1kb bins, Tajima’s D in 1kb bins, and a VCF track.

### Challenges and Prospects for Re-Use of Public Data

Publically available datasets provide opportunities for comparative genomics, but also present limitations inherent to data re-use. Metadata associated with genomic data is often fragmented or missing, meaning crucial information for quality control of reusing data is not always available (Gonçalves and Musen 2019); (Toczydlowski et al. 2021). A key function of the snpArcher pipeline is to produce metrics to evaluate potential biases in the dataset for common population genomic issues. For example, pedigree information is typically not available for wild populations and likely to be missing from public datasets, but close relatives may bias many common population genomic analyses (Hendricks et al. 2018). Our QC module reports relatedness information, allowing rapid identification of related individuals. In the datasets we analyzed, 14% of all datasets considered included identical individuals by genotype and 47% of datasets included at least one first degree relative in it. At the population scale, undetected population structure can bias population and quantitative genomic analysis, and the PCA and admixture reports in the QC module will give a first pass assessment of known or unknown structuring.

Sequencing data on public databases can contain contamination, either from other individuals or other species. These can be identified using measures of inbreeding (i.e., low inbreeding values may suggest excess heterozygosity and cross contamination) that are reported in the QC module. Outliers in sequencing depth, missingness, and mapping rate are all quickly identifiable using the interactive QC plots. Finally, data quality at short scale genomic intervals can be visualized using the genome browser outputs, for example to evaluate sequencing depth and genetic diversity around regions of interest.

### Conclusion

The production of high-quality and accurate genomic variation datasets for non-model species can be a challenging task, especially with the ever-increasing volume of genomic data that is being produced. The massive scale of population-scale whole-genome sequencing datasets presents significant hurdles in data management, processing, and analysis. In this manuscript, we introduce snpArcher, a powerful and user-friendly Snakemake workflow that addresses these challenges and enables the production of reliable and reproducible variation datasets.

Crucially, our pipeline is parallelized, efficient, and scales well even up to modern population-scale datasets. snpArcher also provides an ideal tool for reanalyzing population-level datasets that are available on public databases and provides a consistent framework for comparative analyses across different datasets. By offering a reproducible and well-documented analysis pipeline, snpArcher ensures the reliability and consistency of results, empowering researchers to spend less time on complex data and workflow management, and more time on analysis and discovery.

## Methods

### Workflow overview

The snpArcher pipeline is implemented in Python and utilizes the Snakemake workflow management system (Mölder et al. 2021), which allows for efficient handling of external dependencies, seamless execution on both cloud and cluster computing infrastructures, and enables scalable and efficient analysis of large-scale genomic datasets. For each step in the workflow, snpArcher by default implements tools that are generally regarded as field standard, optimized based on our experience and evaluation of their performance on real-world datasets, ensuring that the pipeline delivers accurate and reliable results. Moreover, snpArcher’s modular and configurable design allows users to customize the analysis workflow to meet their specific research requirements, and allows for future extensions to incorporate additional tools and algorithms.

### Configuration

Core workflow options in snpArcher are controlled by a YAML configuration file. This file controls options such as module selection, output prefix for final files, and temporary storage location. In order to determine what outputs to create, snpArcher requires users to create a *sample sheet file*. This comma separated file contains the required sample metadata about the user’s samples in order to run the workflow. At a minimum, the snpArcher pipeline requires that each sample have a unique sample name, a reference genome accession or a path to a fasta file, and a SRA accession or path to two paired end fastq files (Table 1). We include with snpArcher a simple script, written in Python, to facilitate the generation of sample sheets from local datasets, and we include examples of how to create snpArcher sample sheets from SRA run tables in R.

**Table 1.** An example of the minimum required sample metadata to run snpArcher.

| BioSample | refGenome | Run |
| --- | --- | --- |
| SAMEA3532857 | GCF_003957565.2 | ERR1013161 |
| SAMEA3532860 | GCF_003957565.2 | ERR1013164 |
| SAMEA3532862 | GCF_003957565.2 | ERR1013166 |
| SAMEA3532864 | GCF_003957565.2 | ERR1013168 |
| SAMEA3532865 | GCF_003957565.2 | ERR1013169 |

### Computer resources and cloud configuration

Variant calling for large population-level sequencing datasets are computationally intensive and require substantial computational resources to run. While it is possible to run snpArcher on a laptop for small datasets, such as the test dataset included in the workflow or single samples, we have optimized it to run on HPC clusters and cloud compute platforms. We have tested snpArcher extensively on SLURM-based high-performance clusters and on the Google Cloud Life Sciences platform, and following Snakemake best practices, we provide configurable profiles that can be enabled depending on which computational resources you will use. The SLURM profile and associated bash script provide the basic configuration for running on a SLURM cluster, but the profile will need to be adjusted according to the configuration of the user’s specific system.

To run snpArcher on the Google Cloud Platform, the user must have a Google account linked to a billing account where charges for computational resources can be made. This Google configuration is set up outside of snpArcher, on the command line, and on the Google Cloud Console. Once this is set up in the local environment, snpArcher can be directed to run on Google Cloud instances using the GCP profile provided with the workflow. The user can define how many instances to create and also define the size of required resources in the resources.yaml file included in the workflow. The GCP profile also is configured to exploit preemptible instances, which are short-term compute instances that are offered at considerable cost savings, but can only run for 24 hours and be bought out by other GCP users. The current defaults have been optimized for datasets of genome size of ∼2Gbp, 150 individuals, and 10x sequencing depth with an estimated cost of $1/sample when a Sentieon license is available.

Larger or smaller projects may need to tweak these resources to optimize cost/saving benefits best and prevent the preemption of long-running datasets.

### Data Acquisition and pre-processing

The first step of the workflow is acquisition and pre-processing for raw sequence data and reference genomes. For each sample, two paired-end fastq files are required. The default behavior is to retrieve sequencing data from NCBI based on an SRA run accession (Leinonen et al. 2011) using *prefetch*. For various reasons, *prefetch* may fail. If this happens, *ffq* (Gálvez-Merchán et al. 2022) is used to generate a FTP link for the accession that is downloaded. Alternatively, users can supply paths to fastq files in the sample sheet, in which case snpArcher will operate on those locally stored files. Next, sequencing adapters are trimmed from the raw fastq files with *fastp (Chen et al. 2018)* and sequences with greater than 40% of bases with a phred score below Q15 removed. Reference genomes are retrieved using the NCBI datasets tool (Sayers et al. 2021) if an NCBI accession is specified; otherwise, a path to the reference fasta must be included in the sample sheet. Once available, the reference fasta is processed using *bwa index (Li and Durbin 2009), samtools faidx*, and *samtools dict (Li et al. 2009)* to produce the indexes necessary for downstream processes.

### Read mapping

After the raw data is retrieved and pre-processed, the workflow aligns sequencing reads to the reference genome using *bwa mem (Li 2013)* to produce per sample BAM files. For each sample, read groups are appended based on the sample sheet specification. We mark PCR duplicates using *sambamba markdup (Tarasov et al. 2015)* to exclude these technical artifacts from downstream analysis. Alignment statistics are calculated per sample using *samtools flagstat*.

### Mappability and coverage

Additionally, mappability statistics are computed on the reference genome using *genmap* (Pockrandt et al. 2020). Per site coverage statistics are optionally computed and aggregated using *d4tools (Hou et al. 2021), mosDepth* (Pedersen and Quinlan 2018), and *bedtools (Quinlan and Hall 2010)*. Mappability statistics for the reference genome, combined with per site coverage statistics, can be used to generate a bed file delineating callable regions of the genome based on user-configurable thresholds.

### Variant calling

We use the Genome Analysis Toolkit (GATK) (McKenna et al. 2010) for variant calling and joint genotyping. First, we employ GATK HaplotypeCaller to call SNPs and indels in each sample. If the user has selected the low coverage configuration, we set the *--min-pruning* and *--min-dangling-branch-length* options equal to 1 (Hui et al. 2020), otherwise defaults are used. Next, individual variant calls are aggregated into an efficient data structure via GATK GenomicsDBImport. This step is necessary to enable large cohort joint genotyping. Then, we use GATK GenotypeGVCFs to perform joint genotyping and produce a multisample VCF, retaining only high confidence variants. This approach is broadly adapted by the field as the standard for variant calling, as evidenced by nearly 20,000 citations of the flagship GATK paper to date. Finally, we apply filter annotations to the VCF according to the GATK best practices (Van der Auwera et al. 2013) using GATK VariantFiltration.

### Parallelization

Processing even moderately sized datasets can be exceptionally slow with GATK. One solution is to parallelize each GATK step by splitting the reference genome into processing intervals for both the individual and joint genotyping steps. Optimally, this interval creation step divides the genome into shorter sub-chromosomal (or sub-scaffold) pieces so that each interval can finish in a shorter amount of time. In order to optimize runtime, we use a two-step interval creation process. We generate an initial set of calling intervals using the ScatterIntervalsByNs tool to divide the reference genome at large blocks of Ns. This is important because SNP calling with GATK Haplotype Caller is based on local reassembly, which can be adversely affected if, for example, reads that map across an interval boundary are discarded. However, for many reference genomes this can result in thousands of intervals, which leads to inefficient workflows as the time to assess which jobs need to run becomes prohibitive. To create a balanced set of interval lists, we use the GATK SplitIntervals tool using the option <*-mode BALANCING_WITHOUT_INTERVAL_SUBDIVISION*>, which creates a set of interval lists (up to a maximum user specified value) that all have approximately equal numbers of bases. For the joint genotyping step, each site is treated independently, so we can gain efficiency by creating additional intervals without the concern of splitting adjacent regions of the genome. Thus, for the second set of intervals, we use the option <*-mode INTERVAL_SUBDIVISION*> to produce a scalable number of intervals that can divide adjacent regions. These intervals are then used to parallelize GATK GenomicsDBImport for efficient multi-sample calling.

### Sentieon accelerated variant calling

In addition to the BWA/GATK mapping and variant calling pipeline, we include a Sentieon (Kendig et al. 2019) workflow. This software package is proprietary and produces identical results as GATK, but has been much more efficiently parallelized, resulting in substantially reduced compute needs. The Sentieon workflow uses Sentieon’s drop-in replacement tools for mapping, PCR duplicate removal, metrics, and variant calling. The use of this workflow is a user-specified option in snpArcher and requires a software license from Sentieon that can be specified in the config file.

### Quality Control

snpArcher includes an optional QC module that aggregates various statistics from the workflow and produces preliminary analyses and plots in an interactive HTML file. We estimate the per-individual variant metrics SNP-depth, individual missingness, heterozygosity, and transition/transversions, using *vcftools* v0.1.16 (Danecek et al. 2011). We next generate a small subset of variant data for calculating several preliminary population genomic statistics. In order to generate this pruned dataset, we use *bcftools* v1.12 (Danecek et al. 2021) to first remove all SNPs not passing the filters described above, remove indels, sites with minor allele frequency < 0.01, (i.e., sites present in only 1% of the population), sites with > 75% missing data, and any sites mapping to a previously annotated mitochondrial genome. We next calculate how large of a window to prune this filtered dataset to retain 100k variant sites (i.e. WindowSize = N_SNPs_ / 100,000) and use *bcftools* to select one SNP at random per window. This pruned variant file of 100k SNPs is used for all downstream QC calculations, however, several basic summaries (total number of SNPs, approximate theta, and number of individuals) are calculated from the full variant file and presented in the header of QC HTML file.

We used *Plink2* v2.00a2.3 (Chang et al. 2015; Shaun Purcell)(Galinsky et al. 2016) to perform genome PCA and a KING relatedness matrix (Manichaikul et al. 2010). We also generate a distance matrix using *Plink v* 1.90b6.21 (Purcell et al. 2007). If geographic coordinates are provided, samples will be plotted on an interactive map. Lastly, we used *admixture* v1.3.0 (Alexander et al. 2009) to calculate admixture for *k=2* and *k=3* from the pruned variant file. The output of these analyses, tabulations of variant files, and mapping statistics are all summarized in a single interactive HTML dashboard. Briefly, we use R v4.1.3 (R Core Team 2022) and the following packages for building this summary: *tidyverse* v1.3.1 (Wickham et al. 2019) for data manipulation and *ggplot2* v3.3.5 (Wickham 2016) for graphics, *plotly* v4.9.4.1 (Sievert 2020) for interactive graphics, *ape* v5.5 (Paradis and Schliep 2019) and *ggtree* 3.2.0 (Yu et al. 2018) for phylogenetic tree visualization, reshape2 v1.4.4 (Wickham 2007) for data management, and *ggmap* v3.3.0 (Kahle and Wickham 2013) for terrain maps.

### Postprocessing

In order to enable users to efficiently filter individuals from their VCF file after initially running snpArcher, we include the postprocessing module. Users can trigger this module by marking individuals for removal using the “SampleType’’ column in their sample sheet. The postprocessing module applies customizable filters, which by default remove sites in regions of low mappability and excessive or insufficient coverage (as defined in the configuration file) using *bedtools*, and sites with a minor allele frequency < 0.1 or missingness > 75% using *bcftools* (after recalculating these metrics following sample removal). We also produce separate variant files for SNPs and small indels called by GATK.

### Trackhubs

To display population genomic statistics calculated from the VCF generated by snpArcher, we include an optional module to generate a UCSC Genome Browser track data hub (Raney et al. 2014). At time of publication, this module calculates Tajima’s D (Tajima 1989), SNP density, nucleotide diversity (Pi) and allele frequency. These statistics are calculated using VCFtools v0.1.15 and converted to bigBed format using *bedToBigBed* (Kent et al. 2010).

### Annotating codon degeneracy and inferring synonymous and nonsynonymous variants

snpArcher also includes an optional module that annotates the degeneracy of all coding regions in the reference genome and implements the classic MacDonald–Kreitman test for detecting selection acting in coding regions within a population (McDonald and Kreitman 1991). Briefly, this test compares the number of SNPs present within the population that either change (non-synonymous) or do not change (synonymous) the amino acid encoded at that position. This is compared to similar counts of fixed differences in a diverged outgroup sample to see if and how the ratio of non-synonymous to synonymous changes differs between them. While annotating degeneracy and computing tables for the MK test are common tasks in population genetics, we are not aware of any tools that automate these analyses at a genome-wide scale. To facilitate integration of this functionality into snpArcher, we developed a standalone tool called degenotate (https://github.com/harvardinformatics/degenotate), which calculates MK tables, performs degeneracy annotation, and allows users to extract coding sequences from a genome by their degeneracy.

To implement the MK test across diverse organisms, we make some assumptions about how to classify polymorphic and divergent sites. We consider a polymorphic site to be anywhere at least one ingroup individual has a non-reference allele and fixed differences to be only those sites where none of the outgroup alleles exist in the ingroup. Using these definitions it is possible for a site to both be polymorphic and fixed if the outgroups alleles are different from the alleles segregating within the population. For quantifying variants, we also make some simplifying assumptions. First, if a codon has more than one variant segregating within a population (either because multiple positions at the codon have segregating sites, or because one position has a multi-allelic SNP), we treat each segregating variant as independent. For the outgroup, if there are multiple fixed differences in a single codon in the outgroup, we compute all possible mutational pathways between the ingroup codon and the outgroup codon, and take the average number of nonsynonymous and synonymous changes across these paths, weighted equally. This means we can have fractional numbers of synonymous and nonsynonymous divergence. We also implement calculations of the Neutrality Index (Rand and Kann 1996) and Direction of Selection (Stoletzki and Eyre-Walker 2011) based on the MK test results.

### Empirical datasets

In order to test our pipeline and provide a robust set of consistently processed variant calls for downstream applications, we ran snpArcher on a set of publicly available resequencing datasets (Supplemental Table 1). We focus on non-mammalian vertebrates, as high quality reference genomes are frequently available in this group, but genome sizes are manageable to limit the computational demands needed to process many large population samples. We used SRA to search for possible datasets for inclusion, limiting our search space to species with a) a reference genome, and b) at least one BioProject which contains a minimum of 10 BioSamples sequenced to at least 5x average coverage. The resulting list was then manually curated to identify publications associated with each BioProject, excluding from further consideration datasets for which a publication could not be identified. We then manually assessed the resulting plausible samples to identify a subset for further analysis. R notebooks are provided on Github that contain the code for initial and final assessments (https://github.com/sjswuitchik/compPopGen_ms).

### Benchmarking

To investigate the impact of low sequencing depth on variant calling by, we subsampled the original high-depth dataset Zebra finch dataset to 4x, 10x, and 20x coverage. We ran snpArcher on these subsampled datasets and filtered the resulting VCF files by removing sites not passing standard filters, and calculated heterozygosity statistics using VCFtools v0.1.15 (Danecek et al. 2011). Second, we assessed the effectiveness of our variant calling parallelization (scatter-by-Ns) approach to the conventional (scatter-by-chromosome) approach using the 10x dataset. We performed these benchmarking runs on Google Cloud compute instances, selecting the instance types for each rule to balance cost and runtime (Table S2).

## Data availability

The snpArcher source code is available at https://github.com/harvardinformatics/snpArcher. The Comparative Population Genomics Data public collection is freely available on Globus (https://www.globus.org/). The Zebra finch WGS data used to benchmark snpArcher is publicly available via the SRA BioProject accession PRJEB10586. Scripts used to assess public datasets for the Comparative Population Genomics Data public collection are available at https://github.com/sjswuitchik/compPopGen_ms.

## Acknowledgements

We are grateful for the computational resources provided by the FASRC Cannon cluster, supported by the FAS Division of Science Research Computing Group at Harvard University. CDM, EE, and MB were funded by California Conservation Genomics Project, with funding provided to the University of California by the State of California, State Budget Act of 2019 [UC Award ID RSI-19-690224]. TS was funded by the National Science Foundation (NSF) Division of Biological Infrastructure award DEB-1754397.

